# PROSTATE CELL HETEROGENEITY AND CXCL17 UPREGULATION IN MOUSE STEROID HORMONE IMBALANCE

**DOI:** 10.1101/2024.04.24.590980

**Authors:** Samara V. Silver, Kayah J. Tucker, Renee E Vickman, Nadia A. Lanman, O John Semmes, Nehemiah S. Alvarez, Petra Popovics

## Abstract

Benign prostatic hyperplasia (BPH) is a prevalent age-related condition often characterized by debilitating urinary symptoms. Its etiology is believed to stem from hormonal imbalance, particularly an elevated estradiol-to-testosterone ratio and chronic inflammation. Our previous studies using a mouse steroid hormone imbalance model identified a specific increase in macrophages that migrate and accumulate in the prostate lumen where they differentiate into lipid-laden foam cells in mice implanted with testosterone and estradiol pellets, but not in sham animals. The current study focused on further characterizing the cellular heterogeneity of the prostate in this model as well as identifying the specific transcriptomic signature of the recruited foam cells. Moreover, we aimed to identify the epithelia-derived signals that drive macrophage infiltration and luminal translocation.

Male C57BL/6J mice were implanted with slow-release testosterone and estradiol pellets (T+E2) and harvested the ventral prostates two weeks later for scRNA-seq analysis, or performed sham surgery. We identified *Ear2*+ and *Cd72*+ macrophages that were elevated in response to steroid hormone imbalance, whereas a *Mrc1*+ resident macrophage population did not change. In addition, an Spp1+ foam cell cluster was almost exclusively found in T+E2 mice. Further markers of foam cells were also identified, including *Gpnmb* and *Trem2*, and GPNMB was confirmed as a novel histological marker with immunohistochemistry. Foam cells were also shown to express known pathological factors *Vegf, Tgfb1, Ccl6, Cxcl16* and *Mmp12*. Intriguingly, a screen for chemokines identified the upregulation of epithelial-derived *Cxcl17*, a known monocyte attractant, in T+E2 prostates suggesting that it might be responsible for the elevated macrophage number as well as their translocation to the lumen.

Our study identified macrophage subsets that respond to steroid hormone imbalance as well as further confirmed a potential pathological role of luminal foam cells in the prostate. These results underscore a pathological role of the identified prostate foam cells and suggests CXCL17-mediated macrophage migration as a critical initiating event.

## 1. Introduction

Benign Prostatic Hyperplasia (BPH) is a complex pathological disorder in the prostate mainly characterized by the formation of non-malignant nodules in the transition zone.^1,2^ The prevalence of BPH progressively increases with age affecting 70% of men in their 60s.^3^ Among men aged 50 years and older, BPH is the fifth most common non-malignant disease and represents a significant economic burden in the US.^4^ BPH instigates a range of urinary problems, such as voiding and irritative symptoms, collectively known as lower urinary tract symptoms (LUTS), which can significantly diminish the quality of life.^5^ LUTS/BPH is also linked to a range of complications, including but not limited to urinary tract infection, acute urinary retention, urolithiasis, and renal failure.^6^ Meanwhile, current medical modalities targeting smooth muscle dysfunction via alpha-blockers and proliferation via 5-alpha reductase inhibitors often fail^7^, suggesting that other prostate pathologies play important roles in BPH etiology.

Inflammation and fibrosis are two interconnected processes proposed to drive BPH progression and symptomology. Chronic inflammation is associated with BPH, particularly as the size of the prostate increases, characterized by the dominance of T lymphocytes and macrophages.^8,9^ In turn, chronic inflammation drives fibrosis leading to the worsening of urinary issues due to an impact of urethral resistance.^10,11^ The instigation of chronic inflammation, fibrosis, as well as proliferation in the prostate is thought to be linked to the changing steroid hormone level and the increasing dominance of estradiol with aging.^12,13^ To further decipher this connection, our group has previously utilized a mouse model of steroid hormone imbalance that is created by the subcutaneous implantation of slow-release testosterone and estradiol pellets.^14,15^ We found that macrophages were the primary type of immune cell to be significantly increased in this model, with the ventral prostate lobe experiencing the most substantial rise.^15^ Most importantly, we identified that macrophages migrate to the lumen in both the mouse model and human BPH and take up lipids and adopt the foam cell phenotype.^15^ However, their significance in BPH progression remains largely unclear.

The accumulation of foam cells in the arterial wall is a hallmark of early atherosclerotic lesions, known as fatty streaks.^16^ Furthermore, foam cells undergo programmed cell death, leading to the formation of a necrotic core and inflammatory cell infiltration within the plaque, which is a key feature of advanced atherosclerotic lesions.^16,17^ Foam cells have also been indicated to drive fibrosis in the lung.^18^ These findings imply that foam cells will most likely promote inflammation and fibrosis in the prostate.

To gain a deeper insight into the pathological impact of the shifting immune environment due to steroid hormone imbalance, we utilized single-cell RNA sequencing (scRNA-seq) on the ventral prostate lobe and examined alterations in cell clusters. This has identified a marked increase in progenitor and basal epithelial cells as well as revealed four distinct macrophage subsets, including an *Spp1*+ foam cell cluster. Additionally, we identified the marker gene signatures and pathological factors exhibited by *Spp1*+ macrophages, which will be crucial for future efforts to target these cells and determine their function. Finally, from the transcriptomic changes in resident prostate cells, we identified key cytokines/chemokines that are implicated in remodeling the immune environment.

## 2. Methods

### 2.1 Ethics statement

All experiments were conducted under approved protocols from the Eastern Virginia Medical School Animal Care and Use Committee (ID:22-011) and in accordance with the National Institutes of Health Guide for the Care and Use of Laboratory Animals.

### 2.2 Mice

C57BL/6J male mice were obtained from The Jackson Laboratory. Animals were maintained on a strict 12:12 h light-dark cycle in a temperature- and humidity-controlled facility with water and food provided *ad libitum*. Testosterone (25 mg) and estradiol mixed with cholesterol (2.5 mg and 22.5 mg, respectively) were compressed as pellets and were surgically implanted subcutaneously to 8-week-old mice, as previously.^14,15^ Control mice underwent sham surgery and ventral prostate lobes were collected two weeks post-procedure. Experimental groups involved 8 mice (2/group for scRNA-seq and 6/group for histological tests). The actual count of data points might vary as a result of tissue loss or the removal of low quality samples.

### 2.3 Cell dissociation, library preparation and sequencing

Paired ventral prostate lobes were collected into one tube for each animal in ice-cold PBS and were transferred to 100 ul of 10 mg/mL (in HBSS) *Bacillus licheniformis* protease (Creative Enzymes, Shirley, NY). Tissues were agitated at 6°C at speeds ranging from 600 to 1000 rpm for up to one hour, with intermittent pipetting, until noticeable cell separation was observed. Cells were then passed through a 40 μm mesh strainer and cell viability was assessed on a Countess 3 FL and ranged between 72-78 %. Following the guidelines from the Chromium Next GEM Single Cell manual, we determined the optimal cell loading to be 7000 cells per sample.

Cells were barcoded and libraries were generated for duplicate samples per group using the Chromium Next GEM Single Cell 3⍰ Kit v3.1. Libraries were created from each animal and concentrations were calculated based on Tapestation data (200-1000 bp). Samples were also quantified with Qubit before sequencing to balance the libraries. Paired-end sequencing was conducted on a NextSeq2000 instrument. Run metrics for scRNA-seq of sham and T+E2 mouse prostate samples were calculated by Cell Ranger v7.2.0 (Table S1).

FASTQ files were aligned to *mus musculus* GRCm39 (Ensembl release 111)^19^ cDNA reference for all annotated RNA was prepared using kallisto (v0.46.1) and bustools (v0.39.4).^20^ Transcript counts were aggregated to gene counts using bustools. The count matrix was imported into R (v4.3.2, Bioconductor v3.18.1) and analyzed using Seurat package (v. 5.0.1)^21,22^ and was filtered such that each cell had a minimum of 200 genes and those genes were detected in at least 3 cells. Cells were retained for further analysis if they contained less than 5% mitochondrial reads and 3000 reads/cell. The cold protease method of digestion may have resulted in low cell quality, so strict filtering for mitochondrial reads was included to limit analysis of stressed cells. Using Seurat, data sets were integrated, normalized, scaled (linear model, no regression), and dimensionally reduced using 30 PCs, followed by UMAP to visualize clusters of cells using the top 3000 variable genes and a resolution of 0.8. A second unsupervised clustering was done for fibroblast clusters using the same strategy as above.

Differential gene expression analysis was conducted utilizing the FindMarkers function integrated within the Seurat package, employing the Wilcoxon Rank Sum test for statistical evaluation. Gene Ontology (GO) analysis was performed using the clusterProfiler package and pathways with adjusted p values <0.05 retained.^23^

### 2.4 Immunohistochemistry

Sections were de-paraffinized and hydrated. Antigen retrieval was performed in a decloaking chamber (Biocare Medical, Pacheco, CA, USA) using citrate buffer pH 6.0. Endogenous peroxidases and non-specific binding sites were blocked with Bloxall (Vector Laboratories, Burlingame, CA, USA), rodent block (Biocare Medical) and horse serum (10%). Primary antibodies anti-NR2F6/Ear2 (ab137496, Abcam, Waltham, MA, USA, 1:3000 dilution), anti-GPNMB (ab188222, Abcam, Waltham, MA, USA, 1:1000 dilution) were added overnight. An HRP-conjugated horse anti-rabbit IgG Polymer (Vector Laboratories) was used as secondary for 30 min and the signal was developed using the SignalStain DAB Substrate Kit (Cell Signaling). For IF, anti-CD68 (ab283654, Abcam, Waltham, MA, USA, 1:1000 dilution) and anti-CD72 (AF1279, R&D Systems, Minneapolis, MN, USA, 1:40 dilution) antibodies were coadministered overnight and were labeled with horse anti-rabbit or anti-goat Dylight 488 and 594 (Vector Laboratories, 1:300 dilution) secondary antibodies for 30 min. Alternatively, an Opal kit (Akoya Biosciences, Marlborough, MA, USA) was used with anti-NR2F6/Ear2 (ab137496, Abcam, Waltham, MA, USA, 1:700 dilution) and anti-CD206/Mrc1 (24595T, Cell Signaling, Danvers, MA, USA, 1:500 dilution) primary antibodies.

### 2.5 In situ hybridization

In situ hybridization was performed with RNAscope using probe sets specific for *Ccl6, Cd209a, Col12a1, Cxcl13, Cxcl17, Fbn1, Vegfa*, and *Tgfb*1 with the RNAscope® 2.5 HD Assay-Dual or -Brown detection system (Advanced Cell Diagnostics, Newark, CA, USA). Reaction specificity was tested with a positive control probe and a negative control with only a probe diluent instead of the probe.

### 2.6 Imaging and analysis

Tissues were imaged with a Mantra 2 Quantitative Pathology Workstation (Akoya Biosciences, Marlborough, MA, USA) with a 40x objective. Six representative images were taken per tissue. Area not containing tissue and prostate lumens were removed to obtain total tissue area to normalize positive cell counts that were manually counted. Both experimental groups were represented on each tissue blocks/slides, and slides from the same tissue and timepoint were processed together.

### 2.7 Statistical analysis

Statistical tests were conducted in Graphpad Prism (Graphpad Software, San Diego, CA, USA). For histological analyses, we used two-tailed t-test when the F-test was significant. Otherwise, the statistical difference was determined by Mann–Whitney non-parametric test. P-values above 0.05 were accepted as significant.

## 3. Results

### 3.1 Cell cluster analysis shows an increase in basal and progenitor epithelial cells in T+E2 mice

Single cell libraries were generated for two prostates per experimental group (sham1, sham2, te2-1, te2-2). It is unclear why the sham control samples had fewer genes identified per cell, but may have been related to adjusting to a longer digestion time with cold protease due to the larger T+E2 ventral lobes. Following data filtering, retained cell counts were as follows: 1255 cells for sham1, 1105 for sham2, 1403 for te2-1, and 1031 for te2-2. Using the Seurat R package, data sets were integrated, scaled, and dimensional reduction was performed to find clusters of cells, and genes enriched within each cluster.

Nineteen clusters were first identified (Fig. S1) and canonical lineage-defining markers expressed in the integrated dataset (based on shared genes across treatment groups) were used to annotate clusters (Table S2). Three epithelial clusters deemed highly similar were merged (defined as EpiA cells). Two clusters of prostate basal cells were identified using *Krt5* and *Krt14* markers and subsequently combined. Three presumed epithelial clusters (clusters 7,8, and 18) possessed an overall downregulation in epithelial marker expression potentially indicating low viability in these clusters and were therefore removed from further analysis. A total of five clusters showed marker gene *Krt8* and *Krt18* expression as signatures of luminal cells after merges and the elimination of low-viability clusters (Fig. 1A and B, Table S3). Basal epithelial cells were identified via *Cp, Pdgfa, Apoe* and *Col4a2* expression (Fig. 1B, Table S3). EpiA cells were *Sbp*^high^ and *Spink1*^high^ as well as showed marked expression of *Crabp1* (Fig. 1B), and EpiB cells were identified via the expression of *Nkx3-1, Mmp7, Pbsn* and *Msmb*.^24^ We also identified two epithelial clusters that had high expression of *Mki67* proliferation marker as well as cell cycle-associated genes *Cdca3, Ccnb2, Cdc6* and *Ccne2* (Fig. 1B), whereas gene ontology (GO) pathways were associated with RNA processing and DNA replication (Fig. 1C), suggesting that these clusters consist mainly of proliferating luminal cells (ProlifEpi1 and 2). EpiProgenitor cells presented markers of the urothelial epithelial cell lineage identified by Joseph *et al*. and progenitor markers *Krt4, Krt7, Klf5* and *Slc39a8*.^*25*^ These urothelial epithelial cells were previously identified as prostate luminal cells based on localization in proximal ducts. ^25^ Similar progenitor cells were identified by Guo *et al*. via markers *Tacstd2, Krt4* and *Psca* ^24^ and by Crowley *et al*. via a common proximal marker, *Ppp1r1b*.^26^ EpiProgenitor cells were enriched in metabolic pathways (Fig. 1C). A separate periurethral epithelial cluster that was identified by others via *Ly6d* and *Aqp3*^*26*^ was not found in our dataset.

**Figure 1.**
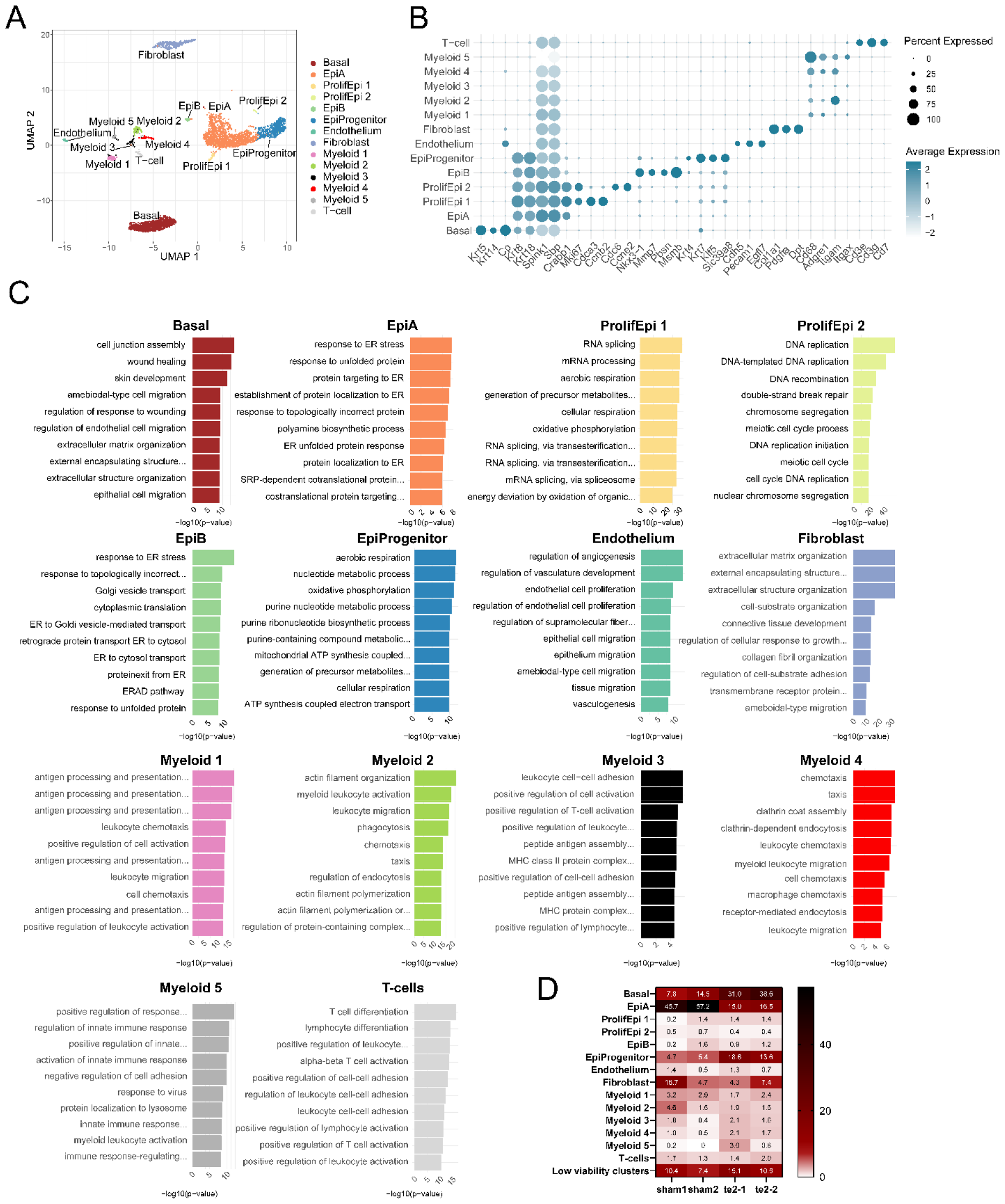
Identification of mouse prostate cell types of sham and T+E2 mice. (A) Major cell lineage clusters based on Uniform Manifold Approximation and Projection (UMAP) embedding of jointly analyzed single-cell transcriptomes from ventral prostate lobes of two sham and two T+E2 mice. After merging highly similar clusters and removing low viability groups (Supp. Fig.) a total of 14 clusters were annotated. (B) Dot blot of selected gene markers used for cell-type identification. (C). Top 10 Gene Ontology terms for each cluster. Pathways with adjusted p values <0.05 were retained. (D) Distribution of cell types within samples shown as % of sample total.

Endothelial cells exhibited positivity for *Cdh5, Pecam1* and *Egfl1* (Fig. 1B, Table S3). Fibroblasts were delineated by the expression of *Col1a1, Pdgfra* and *Dpt*. Myeloid clusters were identified by the expression of *Cd68, Adgre1* (F4/80), *Itgam* and *Itgax*. T-cell identification was based on the expression of *Cd3e* and *Cd3g* and *Cd7*. Smooth muscle cell markers *Acta2* and *Tagln* are expressed in about 5% of cells or less in the fibroblast cluster suggesting that the majority of these cells have been lost during tissue dissociation (data not shown).

Overall cell cluster distribution analysis (Fig. 1D) showed that EpiA cells decreased whereas Basal cells and EpiProgenitor cells were markedly increased in T+E2 mice. In addition, Myeloid 5 cells were almost exclusively in T+E2 mice (2 cells in sham vs. 48 cells in te2). This shows a major rearrangement in cellular heterogeneity in the prostate in response to the two-week exposure to steroid hormone imbalance.

### 3.2 A subepithelial fibroblast cluster is the primary producer of collagens

The fibroblast cluster was further divided into three subclusters via a second unsupervised clustering (Fig. 2A). Marker genes of these fibroblast clusters were identified as *Sult1e1, Igf1* and *Cxcl13* for Fibroblast 1 *Col12a1, Ptn* and *Mfap4* for Fibroblast 2 and *Fbn1, Pla1a* and *Cd55* for Fibroblast 3 (Fig. 2B). The Fibroblast 1 cluster has similar gene signature to what has been identified as “Prostate fibroblast” by Joseph et al.^27^, however marker genes of “urethral” (*Lgr5, Apoe, Osr1* and *Sfrp2*) and “ductal” fibroblast (*Wnt2, Rorb, Wif1, Ifitm1* and *Srd5a2*) were only present at very low levels (data not shown). Extracellular matrix markers of fibrosis, *Col1a1, Col1a2* and *Col3a1* were most pronounced in Fibroblast 3 cells (Fig. 2B). Several GO terms associated with extracellular matrix production were also identified in this cluster (Fig. 2C). These cells might be the main matrix producers in the advanced stages of this mouse model.^15^ Fibroblast 1 cells were linked to increased glucose uptake, whereas Fibroblast 2 cluster was associated with cell-matrix adhesion. Localization of fibroblasts was assessed by RNAscope using *Cxcl13* (Fibroblast 1, red), *Col12a1* (Fibroblast 2, brown) and *Fbn1* (Fibroblast 3, blue) and showed that Fibroblast 1 cells are primarily in the interstitial region whereas Fibroblast 2 and 3 cells are subepithelial (Fig. 2D and 2E). We quantified the expression of this markers in sham vs. T+E2 mice, but did not find significant changes (data not shown), reiterating that fibrosis develops at a later stage in this model.

**Figure 2.**
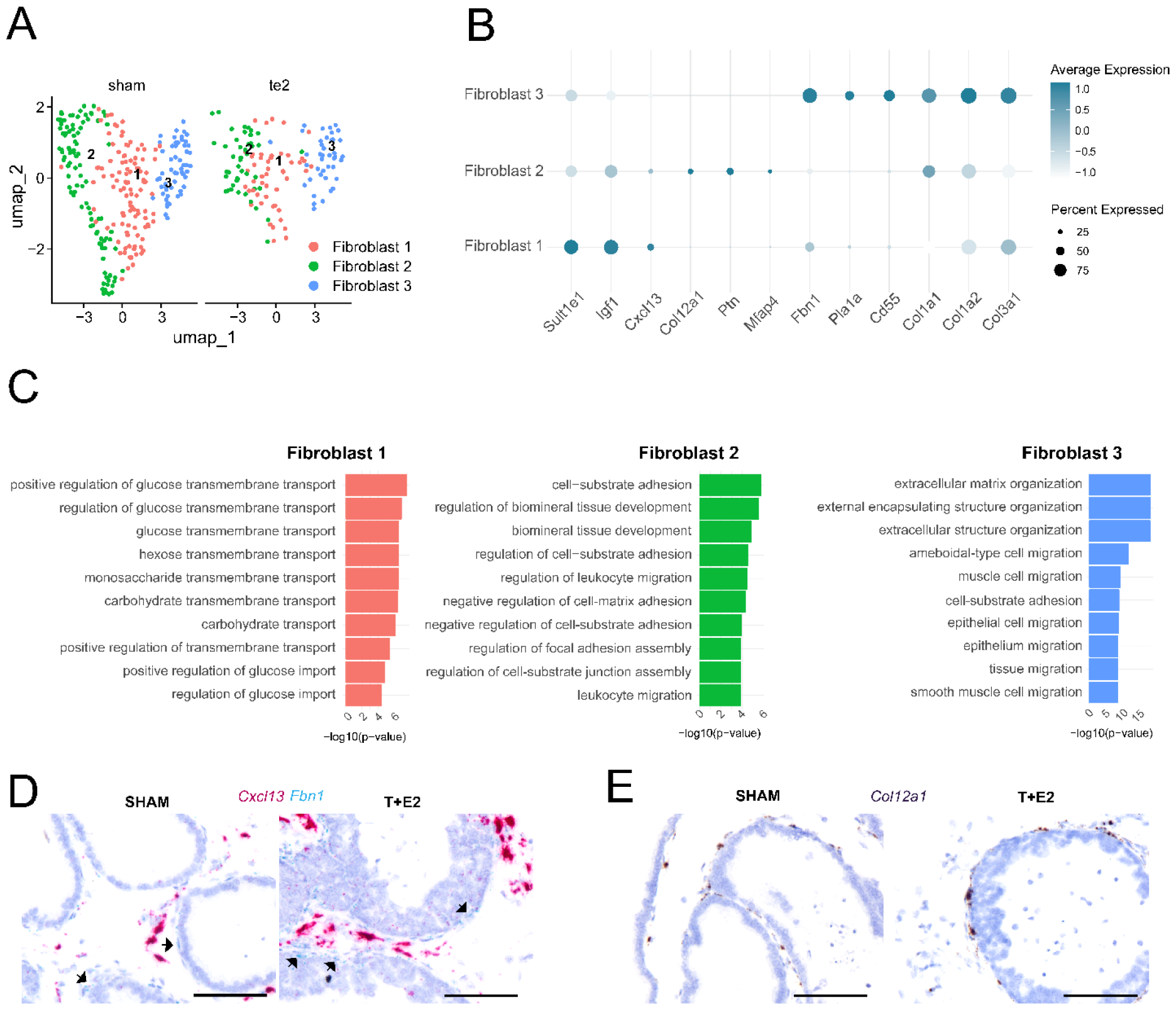
Analysis of fibroblasdt cell clusters in sham and T+E2 ventral prostates. (A) Subclustering of the fibroblast cluster identified 3 fibroblast lineages. (B) Dot blot of markers differentiating the fibroblast clusters (*Sult1e1* – *Cd55*) and delineating extracellular matrix production (*Col1a1, Col1a2, Col3a1*). (C) Top 10 gene ontology terms associated with fibroblast subclusters. Pathways with adjusted p values <0.05 were retained. (D) Localization of Fibroblast 1 (*Cxcl13*^*+*^) and Fibroblast 3 (*Fbn1*^*+*^) cells via *in situ* hybridization. (E) Spatial distribution of Fibroblast 2 cells (*Col12a1*^*+*^) via *in situ* hybridization. Scale bars represent 100 μm.

### 3.3 Intraluminal tissue involution is composed of fibroblasts in T+E2 mice

Histological analysis also identified the presence of incidental intraluminal tissue invaginations that phenotypically resembled stromal nodules and seemed to be rooted in the subepithelial stroma. This is the first time to the best of our knowledge that these structures are described. Calculations of tissue cell counts in our prior analyses excluded these areas. To understand the cell-composition of these structures, we re-assessed fibroblast markers *Col12a1, Cxcl13* and *Fbn1*, as well as E-cadherin epithelial marker in these structures. We found that all three fibroblast markers were expressed in the invaginations (Fig. S2), but not E-cadherin, further confirming the stromal composition of these invaginations. We believe that such structures may be useful in studying the origination of stromal nodules in the future.

### 3.4 Three macrophage clusters increase in response to steroid hormone imbalance

Myeloid clusters were present in all samples, with the exception of the Myeloid 5 cluster, which was almost uniquely composed of cells from the T+E2 samples. (Fig. 3A and B). Using a range of monocyte/macrophage markers based on ^28^, including *Fcgr1, Lyz2, Cd14, Ccr2 and Cx3cr1*, we delineated that clusters Myeloid 1, 2, 4 and 5 are most likely monocyte/macrophage subsets. Interestingly, Myeloid 1 cells expressed a higher level of *Cx3cr1* (Fig. 3C) and *Spn* (Table S3) non-classical monocyte markers, ^29^ whereas Myeloid 2 and 4 possessed elevated *Ccr2* levels representing classical monocytes ^30^ and suggesting that these clusters are composed of specific monocyte lineages and their macrophage derivatives. Genes enriched were *Ms4a7, Cd72, Pmepa1, Fcer1g and Ccl4* in the Myeloid 1 cluster, *Ear2, Fn1, Retnla, S100a6 and Lyz1* in the Myeloid 2 cluster and *Mrc1, Folr2, Mgl2, Gas6, Fcna, Lyve1* and Pf4 in Myeloid 4 cluster (Fig. 3C). The expression of *Folr2* and *Lyve1* suggest that Myeloid 4 cells are resident macrophages whereas *Retnla, Lyz1*, and *Ear2* expression identified the Myeloid 2 cluster as potentially alternatively activated (M2) macrophages.^31,32^ The expression of *Ccl4* and *Cd74* pro-inflammatory markers, and an interferon response gene (*Irf8*) in Myeloid 1 cells (Table S3) suggests a pro-inflammatory phenotype.^33^ In agreement with this, the top GO terms associated with this cluster indicate cell activation and leukocyte chemotaxis (Fig. 1C). Myeloid 3 cells were marked by genes *Traf1, Cd209a, Mcemp1, Rnase6, Wdfy4* and *Btla* delineating these cells as dendritic cells (DCs) most likely derived from monocytes.^31^ Top GO terms in this cluster were related to antigen presentation (MHC complex, T-cell activation, Fig. 1C) in agreement with the proposed cell type function. Most intriguingly, Myeloid 5 cells contained uniquely high levels of *Spp1* (Mac^*Spp1+*^). Our group has previously identified *Spp1* as a marker of lipid-laden luminal macrophages in the mouse ventral prostate otherwise known as foam cells. Examples of genes further distinguishing the Mac^*Spp1+*^ cluster were *Fabp5, Ctsl, Gpnmb, Mmp12* and *Trem2*, which markers are also predominant in foamy macrophages in mouse atherosclerotic lesions.^34^

**Figure 3.**
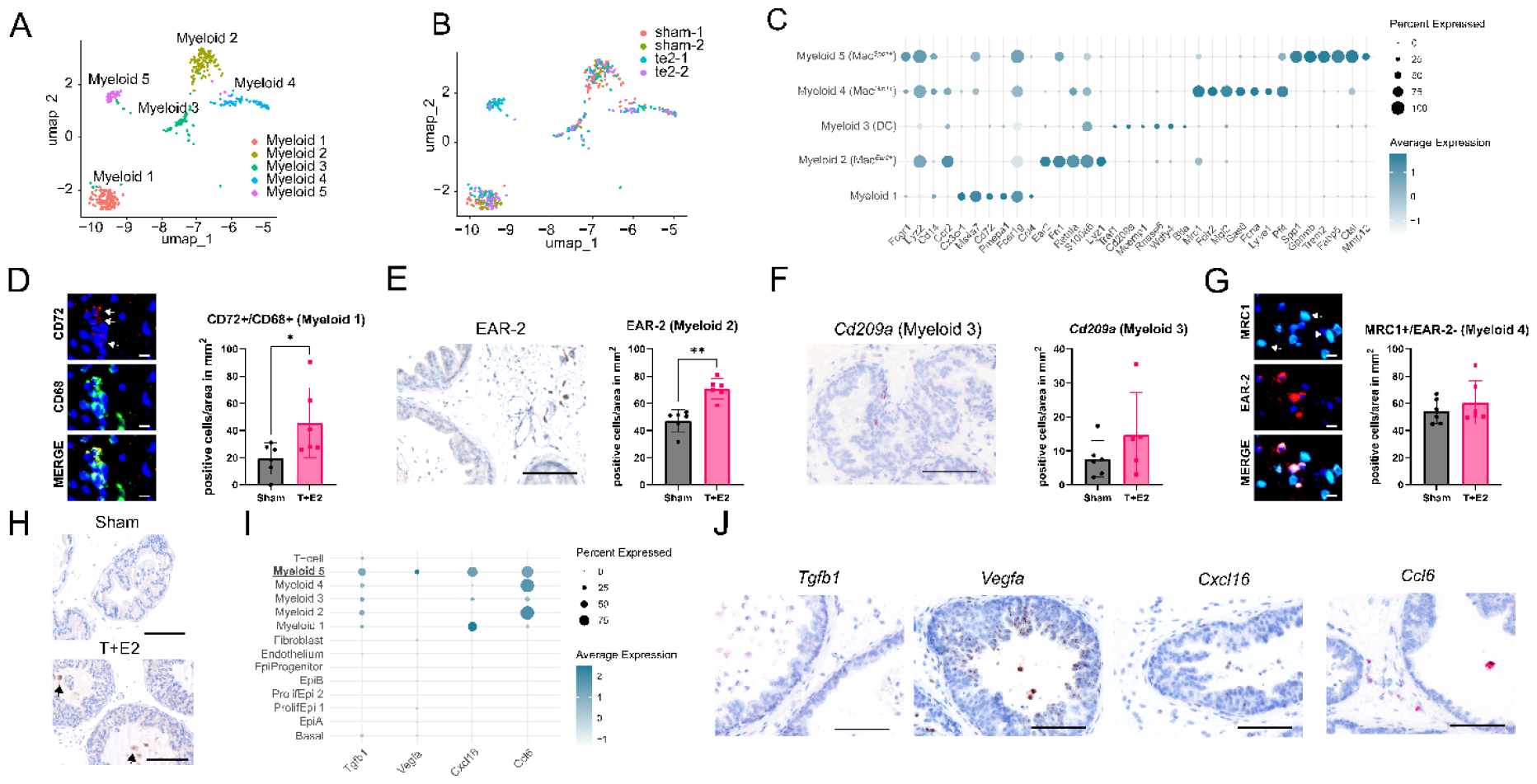
Analysis of myeloid cluster markers and Mac^*Spp1*+^ pathological gene transcriptome. (A) Uniform manifold approximation and projection (UMAP) visualization of myeloid clusters. (B) Distribution of treatment groups across myeloid clusters. (C) Expression of known markers of macrophages/monocytes (*Fcgr1*-*Ccr2*) and our markers of Myeloid 1 (*Ms4a7*-*Ccl4*), 2 (*Ear2*-*S100a6*), 3 (*Traf1*-*Btla*), 4 (*Mrc1*-*Pf4*), and 5 (*Spp1*-*Mmp12*) clusters. (D) Representative image of CD72+/CD68+ immunofluorescence (IF) to identify Myeloid 1 cells and quantification in Sham vs. T+E2. (E) Representative image of EAR-2+ immunohistochemistry (IHC) to identify Myeloid 2 cells and quantification in sham vs. T+E2. (F) Representative image of Cd209a mRNA via *in situ* hybridization to identify Myeloid 3/Dendritic cells and quantification in sham vs. T+E2. (G) Representative image of MRC1+/EAR-2-IF to identify Myeloid 4 cells and quantification in sham vs. T+E2. (H) Representative image of GPNMB of Myeloid 5 luminal macrophages in sham and T+E2 tissue. (I) Further profibrotic, proinflammatory and pro-angioneic factors expressed in Mac^Spp1+^ macrophages. (J) Validation of mRNA expression of *Tgfb1* (red), *Vegfa* (brown), *Cxcl16* (brown), and *Ccl6* (red) in luminal macrophages via *in situ* hybridization. Only examples of T+E2 tissues are shown. Scale bars represent 100 μm except (D) and (G), where scale bars are 10 μm.

Myeloid 1 cells were labeled based on CD72^+^CD68^+^ dual positivity in the tissues and were shown to be increased by T+E2 treatment (Fig. 3D). The addition of the CD68 was required to exclude potential B-cells that are known to be positive for CD72. The EAR-2 positive Myeloid 2 (Mac^*Ear2*+^) macrophages localized to the interstitial stroma and were increased by 50% in response to T+E2 treatment (Fig. 3E). DCs were localized in the periglandular area shown via Cd209a *in situ* hybridization, but were not significantly elevated in response to T+E2 treatment (Fig. 3F). Myeloid 4 cells were labelled using MRC1 (Mac^*Mrc1*+^), however, a small subset of Mac^*Ear2*+^ were also expected to stain for this marker based on Fig. 3C. Therefore, we identified these cells based on MRC1+EAR2-staining and found that they were localized in the stroma and around the epithelium and capillaries, but did not significantly change in number in response to treatment (Fig. 3G). Using immunohistochemistry, we confirmed that GPNMB protein expression is limited to luminal cells with foam cell morphology and were only detected in the T+E2 group (Figure 3H).

### 3.5 *Spp1+* macrophages express *Tgfb1, Mmp12, Cxcl16, Ccl6* and *Vegfa* potential pathological factors

To understand the functional role of the Mac^*Spp1*^ macrophages, we screened their transcriptome for genes that encode secreted proteins and therefore may influence nearby epithelial cells. Mac^*Spp1*^ marker genes with p value < 0.001 were loaded to a Uniprot batch analysis. This search identified 61 genes that encode secreted proteins (Table S7). From these genes, we selected factors with previously identified pathological roles including *Transforming growth factor beta 1* (*Tgfb1*),^52,53^ *Matrix metalloproteinase 12* (*Mmp12*),^58^ *chemokine (c-c motif) ligand 6* (*Ccl6*)^57^ and *Vascular endothelial growth factor-a* (*Vegfa*)^54^ for further analysis via dot blot across cell clusters. We also included *Chemokine (c-x-c motif) ligand 16* (*Cxcl16*), which is primarily categorized as transmembrane protein in the Uniprot database, but has been shown to be a soluble attractant for T-cells.^35^ Dot blot analysis showed that *Cxcl16* was also expressed in Mac2 and *Ccl6* in Mac1 cells (Figure 3I). We then confirmed the expression of these markers in luminal macrophages using RNAscope (Fig. 3J). *Vegfa* showed high expression in luminal macrophages and was also moderately elevated in epithelial cells (Fig. 4C). *Cxcl16, Tgfb1* and *Ccl6* also showed positive staining in luminal cells. In addition, Mac^*Spp1*^ cells in tissue have a foam cell phenotype with elevated intracellular lipid.^15^ Thus, a panel of genes involved in lipid metabolism were evaluated. Mac^*Spp1*^ cells have marked increase in the expression of genes related to lipid uptake (*Cd36*),^36^ intracellular trafficking (*Fabp4* and *Fabp5*)^37^ and efflux (*Abcg1* and *Abca1*, Fig. S3)^38^.

**Figure 4.**
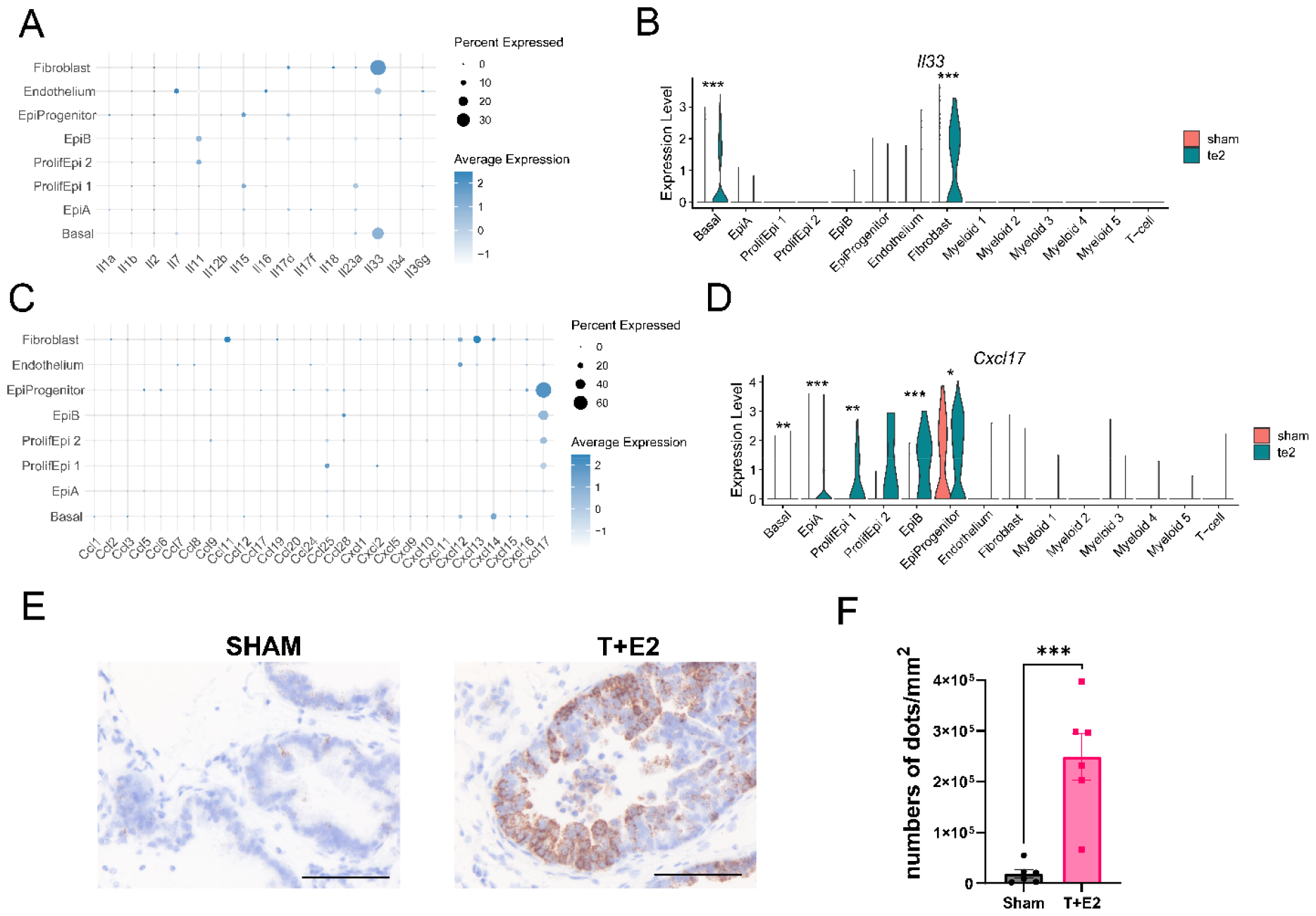
Steroid hormone imbalance stimulates *Cxcl17* expression in epithelial cells. (A) Dot blot showing the expression of interleukines in resident prostate cells. (B) Violin plot demonstrating expressional changes in *Il33* in control and T+E2 ventral prostate in all cell clusters. (C) Dot blot showing the expression of chemokines in resident prostate cells. (D) Violin plot demonstrating expressional changes in *Cxcl17* in control and T+E2 ventral prostate in all cell clusters. (E) Spatial localization and expression of *Cxcl17* in sham and T+E2 ventral prostates with *in situ* hybridization. (F) Quantification of *Cxcl17* expression. Scale bars represent 100 μm. Significance was tested using Student’s t-test. ***p⍰≤⍰10.001.

### 3.6 Steroid hormone imbalance upregulates the monocyte attractant *Cxcl17* in epithelial cells

Our previous study,^15^ as well as the current analysis, highlighted that steroid hormone imbalance drives monocyte infiltration to the prostate as well as the translocation of macrophages to the prostate lumen where they differentiate into foam cells. In order to assess how resident prostate cells drive remodeling of the immune environment, we visualized the cytokine expression profile across epithelial, fibroblast and endothelial clusters in T+E2 treated versus control cells (Fig. 4). Differential expression (DE) analysis was completed and statistical analysis was performed using non-parametric Wilcoxon Rank Sum test (Table S6). A screen through all interleukins (ILs) identified *Il33* as being predominantly upregulated in basal epithelial cells and fibroblasts (p<0.001, Fig. 4A,) and induced by T+E2 treatment (Fig.4B). *Il33* is a nuclear cytokine and only activates immunological response upon cell damage and extracellular release.^39^ We also screened chemokine (C-C motif) ligand (*Ccl*) and Chemokine (C-X-C motif) ligand (*Cxcl*) gene expression levels across different cell clusters and then compared selected candidates between sham and T+E2 animals (Fig. 4C and D). This screen identified *Cxcl17* as an upregulated gene in epithelial clusters (Basal p<0.01, EpiA p<0.001, ProlifEpi1 p<0.01, EpiB p<0.001, and EpiProgenitor p<0.05). RNAscope determined a significant 13-fold upregulation in *Cxcl17* expression in T+E2 mice (Fig. 5E and F) and thereby confirmed the scRNA-seq results. CXCL17 is a known monocyte attractant^40^ indicating that it may be the primary driver of the increase in macrophages in the tissue as well as macrophage translocation to the lumen.

## 4. Discussion

In a recent study, we discovered the accumulation of luminal foam cells specifically in the ventral prostate lobe in a mouse model recapitulating human age-related steroid hormone imbalance and in human BPH.^15^ Building on these findings, our current study explores the changing immune cell milieu, the transcriptome of luminal foam cells and identifies secreted factors of resident prostate cells with the potential to drive immune cell infiltration in response to testosterone and estradiol slow-release pellets.

Multiple epithelial cells were observed, including secretory epithelial cell types EpiA and EpiB, consistent with previous scRNA-seq analyses.^24-26^ The representation of EpiB cells was found to be limited, corroborating earlier findings which identified these cells predominantly in the anterior, lateral, and dorsal lobes.^24^ Additionally, our findings highlighted a significant increase in the representation of progenitor “urothelial” epithelial cells ^24-26^ as well as basal cells in response to steroid hormone imbalance. Urothelial cells in the prostate are known to express various progenitor markers and have been characterized as castration-resistant/androgen-independent luminal cells that proliferate in response to androgen-induced stromal growth factors.^25,41^ The analogous cell type in humans is observed to be enriched in BPH patients treated with 5-alpha-reductase inhibitors (5ARIs).^25^ Our study demonstrated that estradiol/testosterone supplementation also produces an increase in urothelial epithelial cells.

Our characterization of prostatic fibroblasts provided three subtypes. Our Fibroblast 1 cluster is identical to “Prostate fibroblast” earlier recognized by Joseph et al.^27^ and showed a highly metabolically active state represented by increased monosaccharide and carbohydrate import pathways. Fibroblast 2 and 3 were mainly localized to the subepithelial region and may potentially be subtypes of the “ductal fibroblasts” identified by Joseph *et al*.^27^ Most importantly, we found the highest expression of collagen genes in Fibroblast 3 cells emphasizing that this cell type could be a driver of prostatic fibrosis in the advanced stages of this model.^15^ Our future studies will confirm this hypothesis in more chronic stages of the model. The lack of smooth muscle cells in our dataset is potentially due to the digestion protocol selected (cold protease), which will be adjusted in the future to retain these cells.

The immune environment captured included four macrophage clusters, one dendritic cell cluster and one T-cell cluster. Based on gene expression signatures, Mac^CD72+^ are pro-inflammatory (M1), Mac^Ear2+^ are alternatively activated (M2), Mac^Mrc1+^ are resident macrophages, and Mac^Spp1+^ are foam cells. Within the resident macrophage cluster, the presence of cells expressing the classical monocyte marker *Ccr2+* suggests that these cells are subject to monocyte-mediated replenishment. Resident macrophages support tissue functionality; Mac^Mrc1+^ cells exhibit *Cd36* levels comparable to those of foam cells, indicating a potential role in lipid homeostasis. However, steroid hormone imbalance led to an increase in all macrophage subsets, except for resident macrophages, indicating the pivotal role of macrophage infiltration in this model. Mac^CD72+^ in the heart were shown to be pro-inflammatory and promote cardiac injury.^42^ Cells with similar marker gene signature have been identified as interstitial macrophages that respond to LPS treatment^43^ and may represent the pro-inflammatory macrophage subset we identified in an earlier study based on iNOS expression.^15^ Conversely, Mac^Ear2+^ (myeloid 2) cells carry Retnla universal and Ear2 monocyte-derived alternatively activated (M2) macrophage markers.^31,32^

Mac^*Spp1*+^ cells were delineated in our prior study as intraluminal foamy macrophages in this model, setting the stage for the goal of our current study to further elucidate their characteristics.^15^ We found that, *Gpnmb* and *Trem2*, recognized foam cell markers in atherosclerosis, were also present in this macrophage subset.^34^ Although foam cells in atherosclerosis are known to exacerbate inflammation through their accumulation and formation of a necrotic core via programmed cell death^16^ their phenotype is closer to an anti-inflammatory M2 state based on markers *Cd9, Ctsb, Fabp4*, and *Lgals3*. ^47-49^ Foam cells may also constitute a heterogenous population where a *Spp1*+*Trem2*-subtype undergoing endoplasmic reticulum stress, apoptosis and autophagy possesses amplified glycolytic and angiogenic activities.^50^ The presence of oxidized lipids induces the expression of pro-inflammatory genes *TNF* and *IL6* in human macrophages, indicating that foam cells might initially present an M1 phenotype.^51^

Our focus was on uncovering the pathological significance of foam cells in the prostate by analyzing their marker gene expression, and identified *Tgfb1, Vegfa, Cxcl16, Ccl6* and *Mmp12* as potential mediators. TGF-β1 is widely recognized as a key factor that promotes fibrosis and concurrently compromises the integrity of the barrier function in prostate epithelial cells.^52,53^ VEGF-A has been implicated in the enhancement of microvascular density within BPH nodules, signifying its contribution to the progression of BPH.^54^ CXCL16 attracts mainly T-cells and promotes fibrosis^55,56^ whereas CCL6 attracts monocytes and macrophages^57^ signifying the potential role of foam cells in initiating an inflammatory response in the prostate. Moreover, MMP12 emerges as a crucial foam cell-secreted enzyme that not only facilitates macrophage migration — thereby influencing inflammatory responses — but also plays a significant role in increasing the permeability of epithelial barriers.^58^ Finally, osteopontin (*Spp1*), the first foam cell-secreted factor in BPH identified in our previous study, is a pro-inflammatory, pro-fibrotic protein.^15,59-61^ Osteopontin is efficiently taken up by adjacent epithelial cells, as demonstrated by immunohistochemistry, highlighting the potential impact of foam cell-derived proteins on tissue homeostasis and disease progression.^15^ Such proteins, especially in areas where epithelial integrity is already compromised, possibly due to the actions of TGF-β1 or MMP12, might indeed augment disease processes. Our future studies will focus on understanding the mechanisms by which these factors contribute to disease to provide valuable insights into potential therapeutic targets for intervention.

We also explored the cytokines involved in monocyte infiltration and macrophage luminal translocation. Importantly, *Cxcl17*, a gene that encodes a secreted chemokine, showed increased levels in luminal epithelial cells in response to steroid hormone imbalance. Past studies have shown CXCL17 to act as a chemoattractant for monocytes/macrophages, exerting a comparatively weaker effect on dendritic cells.^40^ The epithelial expression and potential luminal secretion of CXCL17 also align with expectations for a cytokine that could facilitate the movement of macrophages into the luminal space. Therefore, our future studies will aim to elucidate whether the observed upregulation of CXCL17 in steroid hormone imbalance, mediates macrophage mobilization and infiltration into the luminal compartment, thereby facilitating foam cell formation. Additionally, we found that *Il33* was upregulated in basal epithelium and fibroblasts. Typically, IL33 protein is not secreted and retained in the nucleus, becoming active only when released due to cell damage.^62^ However, more recent findings have shown that IL33 may be secreted through the neutral sphingomyelinase 2 endosome pathway.^63^ IL33 promotes the proliferation and survival of immune cells and triggers a type 2 immune response ^64,65^ but, as discussed above, may only contribute to a more extensive response in the prostate following cell death, such as in inflammatory atrophy.

Within our framework, the direct influence of estradiol on macrophages is also expected to play a role. Estradiol has been shown to diminish LPS-induced NF-κB activation and IL-6 production in monocytes and macrophages. ^66,67^ Furthermore, it reduces monocyte adhesion through the downregulation of Rac1 GTPase activity.^68^ Estradiol also promotes wound healing by inducing M2 polarization through macrophage-specific estrogen receptor alpha (ERα) signaling^69^ collectively underscoring its antiinflammatory properties. The effects of estradiol, however, exhibit sexual dimorphism; it inhibits LDL-induced lipid accumulation in macrophages derived from female subjects but does not affect those from male subjects. ^70^ Estradiol also diminishes M2a activation in macrophages from males. ^71^ However, considering the complexity of an *in viv*o system, the direct effects of estradiol might be inferior to other paracrine factors on macrophages; the direct action of estradiol, for example, should inhibit foam cell formation by decreasing lipid accumulation in macrophages, as previously shown,^72^ which contrasts with our observations in the male steroid hormone imbalance model. Another possibility is that local estrogen levels are highly variable in the tissue microenvironment, especially, when we compare tissue vs. intraluminal areas. Our future studies will aim to test these hypotheses to provide a molecular mechanism of how steroid hormone imbalance drives foam cell formation in the prostate.

A weakness of our study is that the sham control samples had fewer genes identified per cell, which may have been due to the cold protease digestion methodology, but what was kept consistent between the two groups. To address this, we have confirmed all major findings with a secondary method of in situ hybridization or immunohistochemistry. Current work in the lab is ongoing to optimize digestion strategies and to increase sample number to further identify differences in cell populations within the steroid hormone model.

In conclusion, we have characterized changes in prostate cell heterogeneity in the mouse steroid hormone imbalance model in the ventral prostate. We found increases in progenitor and basal epithelial cells as well as in three subsets of macrophages. Most importantly, we have identified important secreted factors specific for luminal foam cells as well as identified CXCL17 as a candidate for directing macrophage trafficking in the prostate. Our future studies will mechanistically decipher the role of these newly identified molecular drivers, especially how luminally-secreted factors drive disease pathogenesis in BPH.

## Acknowledgements

We thank Xin (Cindy) Guo, Chunghwan Ro, Daniel McWilliams, Mary Ann Clements and Dita Julianingsih for experimental assistance and advice. This research was funded by grants from the National Institutes of Health (K01 DK127150 to P.P and 1 U2C CA271894-01 to OJS), financial support from the Hampton Roads Biomedical Research Consortium and a start-up fund from the Eastern Virginia Medical School (to P.P.). The authors thank the EVMS Histology Services Lab for providing its services.

## Author contribution statement

PP and NSA conceptualized the study. PP and OJS acquired funding. PP and NSA developed the methodology. SVS, PP and KJT performed the experiments and conducted the formal analysis. PP and SVS visualized the data. PP wrote the manuscript and REV, NAL, NSA and OJS conducted manuscript review and editing. All authors have read and agreed to the published version of the manuscript.

## Data availability statement

Expression data is accessible through GEO Series accession number GSE263790

