## Supplemental document for "PROSTATE CELL HETEROGENEITY AND CXCL17 UPREGULATION IN MOUSE STEROID HORMONE IMBALANCE"

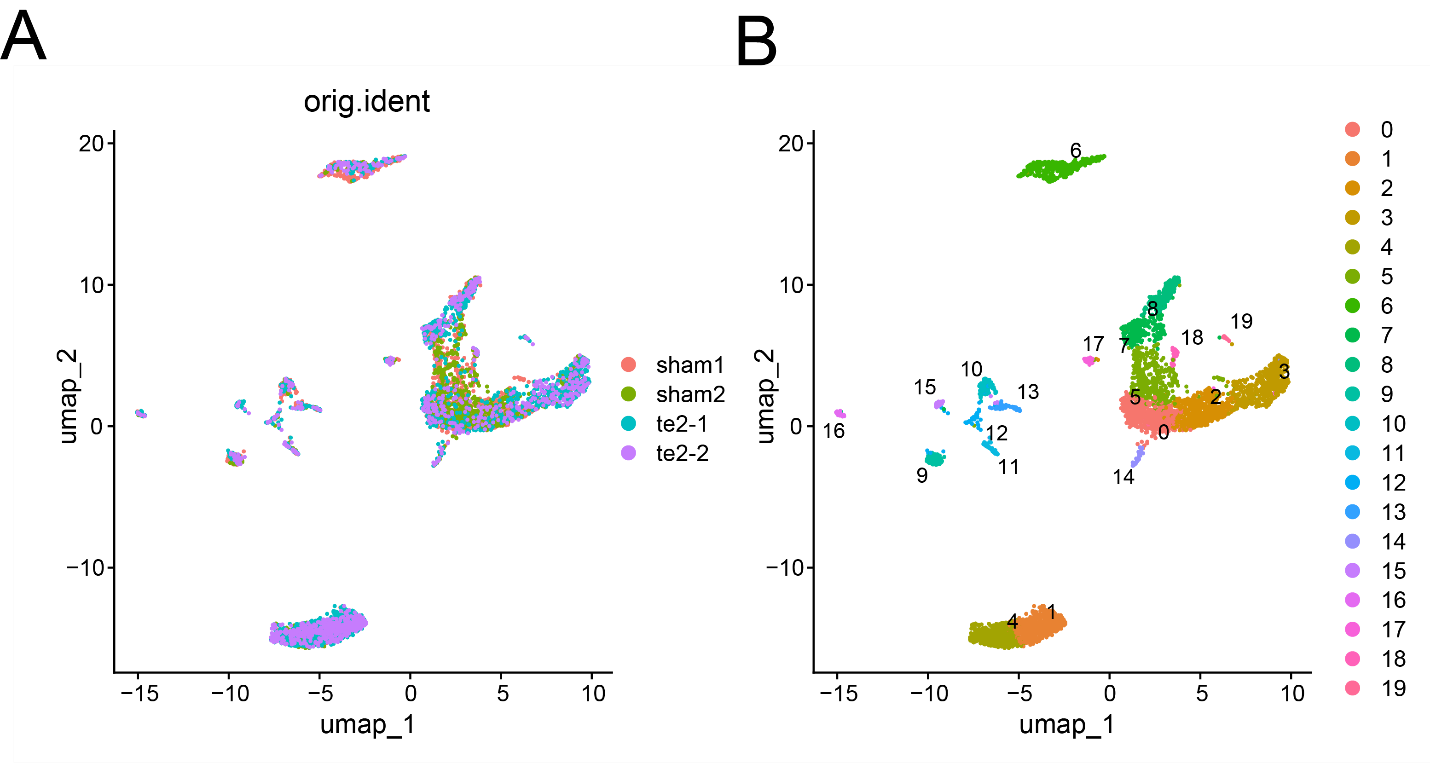
Figure S1: Uniform manifold approximation and projection plot (Umap) of the integrated dataset composed of cells of two sham and two T+E2 ventral prostates. (A) samples were overlayed to demonstrate integration efficiency. (B) Using 3000 Features and a resolution of 0.8, 19 clusters were delineated by the Seurat workflow.

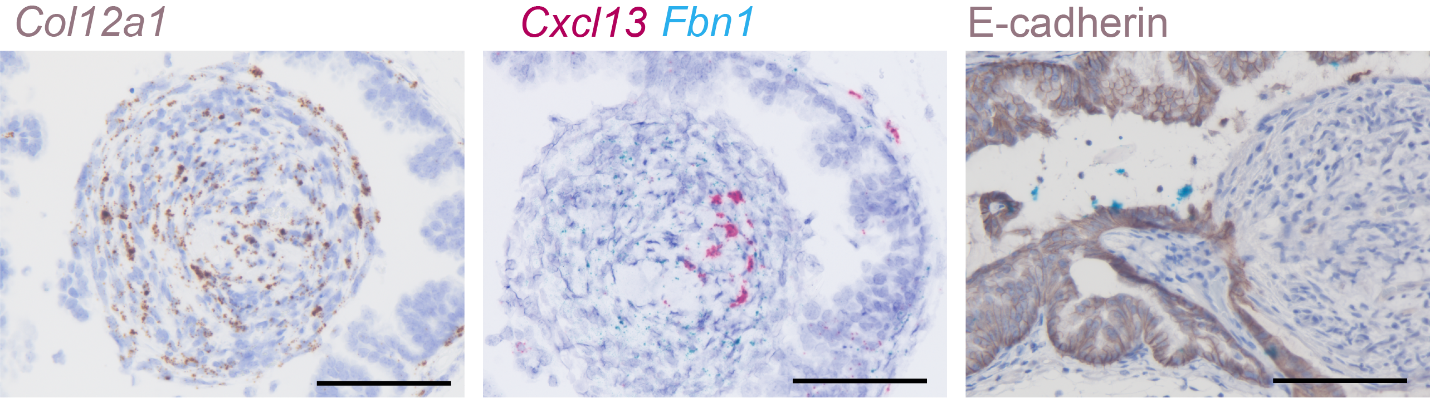

Figure S2.: Cellular composition of intraluminal tissue invagination. *Col12a1*, *Cxcl13* and *Fbn1* fibroblast markers were detected by RNAscope. E-cadherin was labeled with immunohistochemistry. Scale represents 100 µm.

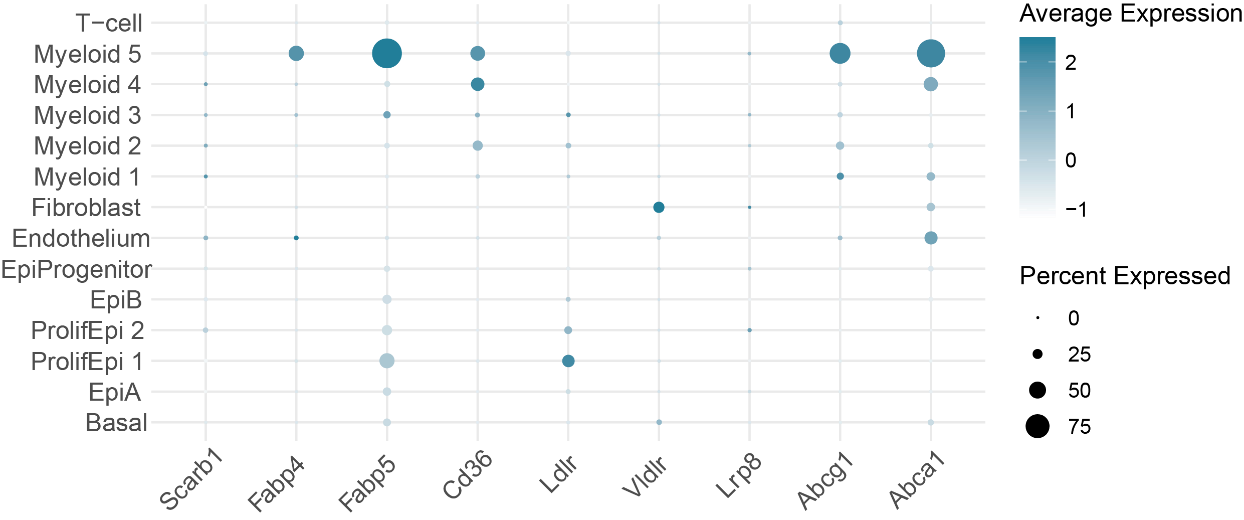

Figure S3: Dot blot expression analysis of lipid transporters across all cell clusters.
