## Supplemental Table 1 for "PROSTATE CELL HETEROGENEITY AND CXCL17 UPREGULATION IN MOUSE STEROID HORMONE IMBALANCE"

**Supplemental Table 1. Run metrics for scRNA-seq of sham and T+E2 mouse prostates.**

| Sample | Number of Reads | % Q30 Bases in Read | % Reads Mapped | Estimated Number of Cells | Mean Reads per cell | Median Genes per cell |
| --- | --- | --- | --- | --- | --- | --- |
| sham1 | 225,394,469 | 94.6 | 88.6 | 2,210 | 101,988 | 619 |
| sham2 | 197,536,680 | 94.4 | 88 | 2,796 | 70,650 | 750 |
| te2-1 | 160,431,243 | 94.4 | 93 | 2,815 | 56,992 | 1,935 |
| te2-2 | 182,727,898 | 94.7 | 93.4 | 2,711 | 67,402 | 2,066 |
